# Cytoplasmic dynamics are overlooked in single nuclei RNA-seq but can be rescued by CytoRescue, a generative AI model to recover cytoplasm enriched gene

**DOI:** 10.1101/2025.08.15.670239

**Authors:** Wankun Deng, Zhongming Zhao

**Author notes:** to whom the correspondence should be addressed: Zhongming Zhao, Ph.D., Center for Precision Health, McWilliams School of Biomedical Informatics, The University of Texas Health Science Center at Houston, 7000 Fannin St. Suite 600, Houston, TX 77030.

## Abstract

Single-nucleus RNA sequencing (snRNA-seq) generates single cell data from nuclei. It provides valuable compatibility with frozen or difficult-to-dissociate tissues while avoiding stress responses in fresh samples. However, cytoplasmic depletion inherently limits quantification of cytoplasm-enriched genes. Here, we present CytoRescue, a novel generative AI model designed to recover attenuated cytoplasmic signals in snRNA-seq data. Our results demonstrate that CytoRescue effectively restores expression of cytoplasm-enriched genes while preserving underlying gene expression signatures. Taking advantaging of the raw-in-raw-out design, CytoRescue can be easily integrated into the existing pipelines for single-cell sequencing analysis. Notably, CytoRescue successfully recovers EGF signaling pathway components, a critical cell-cell communication pathway in lung cancer, in an independent dataset. CytoRescue addresses a fundamental limitation of snRNA-seq technology, enhancing its utility for comprehensive transcriptomic profiling while maintaining the advantages of single nucleus-based approaches.

## Background

Single-nucleus RNA sequencing (snRNA-seq) is considered more flexible with sample preparation when comparing to single cell RNA-seq (scRNA-seq), which requires live cells from fresh tissue followed by quick dissociation in most cases^1^. Benefitting from the compatibility of frozen sample, snRNA-seq is widely used in single cell research^1^. Meanwhile, in snRNA-seq experiments, cytoplasm content will be discarded during nuclei isolation^1^ (Figure 1A), resulting in missingness of cytoplasm dynamic^1–4^. However, various cytoplasm processes exhibit high dynamic in certain cells while related genes showing no or small variation in nuclei^1–4^. For example, microglia activation related genes are enriched in cytoplasm and snRNA-seq cannot accurately detect microglia activation^4^. Furthermore, dynamic in MHC-1 signaling pathway, which is crucial in immune recognition and response, can barely be detected in snRNA-seq in neither normal nor tumor lung samples^3^.

**Figure 1.**
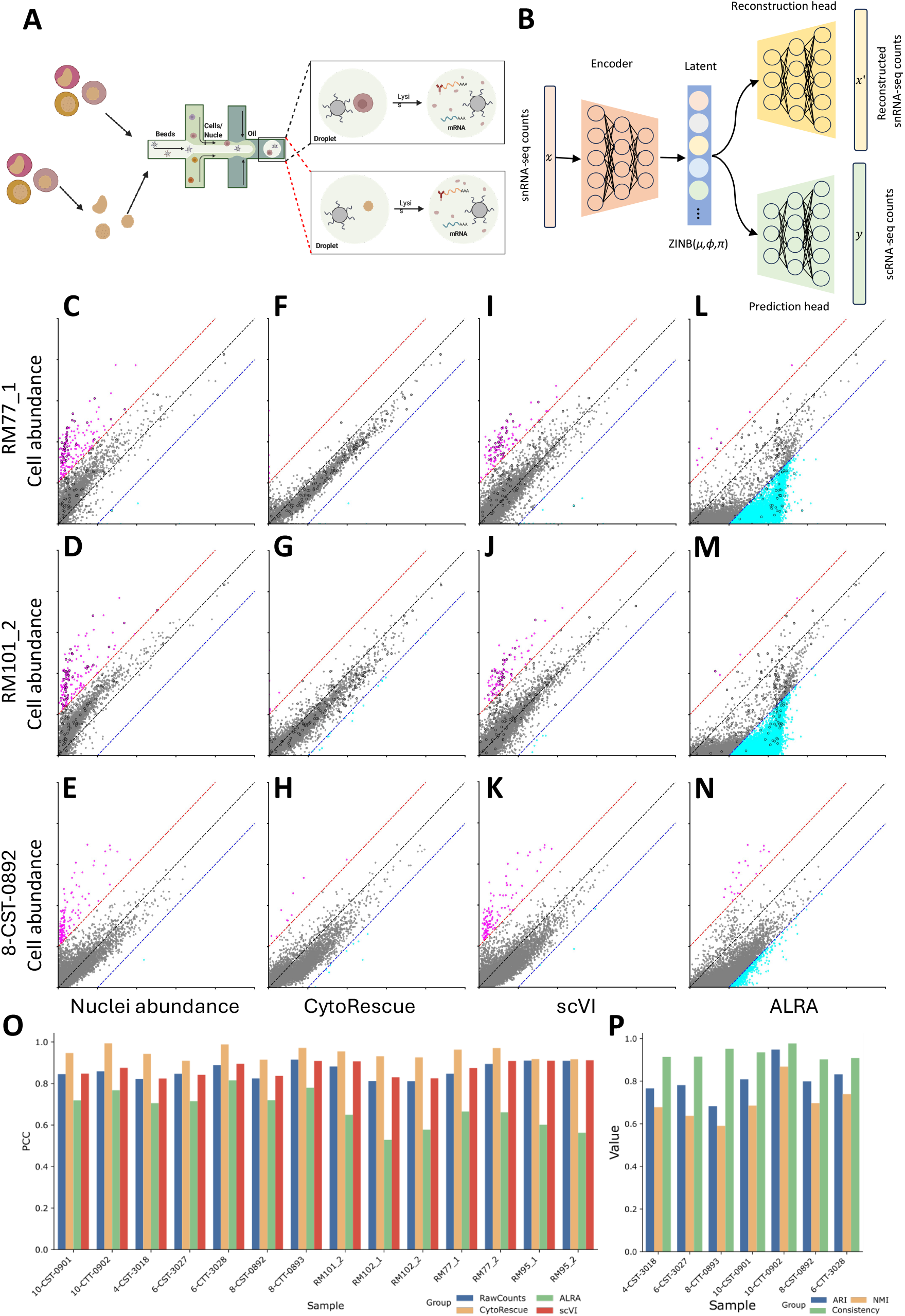
Diagram and evaluation of CytoRescue. (A) An overview of scRNA-seq and snRNA-seq procedures. (B) Design of CytoRescue. The model is designed with both reconstruction and prediction heads. Using paired snRNA-seq and scRNA-seq profiles as prediction targets, CytoRescue is trained on a combined loss function by incorporating reconstruction loss from the reconstruction head, Kullback-Leibler divergence of the latent space, and mean squared error from the prediction head. (C-N) Mean gene expression comparison between scRNA-seq and snRNA-seq by three methods: CytoRescue-predicted, scVI-imputed, and ALRA-imputed profiles. Results from three samples are shown: RM77_1 (brain, training sample), RM101_2 (brain, validation sample), and 8-CST-0892 (normal lung, validation sample). Disease-associated microglia genes are marked as dots with black edges in panels C, D, F, G, I, J, L, M. (O) Pearson correlation coefficients (PCC) between mean gene expression of scRNA-seq profiles and raw snRNA-seq counts, CytoRescue-predicted, scVI-imputed, and ALRA-imputed profiles across all samples in this study. (P) Adjusted Rand index (ARI), normalized mutual information (NMI), and consistency of cell type annotation in lung samples.

The recent advances in generative artificial intelligence (AI) models shed lights on modeling biological structures from biased, noisy and sparse data^5–8^. Among many algorithms, variational autoencoder (VAE) has emerged as one of most popular frameworks in single-cell study, benefitting from its capability to model the probabilistic distribution of single cell read counts^6,8–10^. To model single-cell RNA-seq data, a zero inflated negative binomial (ZINB) distribution was proposed by adding an additional parameter π as zero-inflating probability^11^. By leveraging these progresses, we propose a VAE algorithm based on ZINB distribution, namely, CytoRescue, to rescue the lost cytoplasm dynamic in snRNA-seq experiments due to technology limitation.

## Results and discussion

We developed CytoRescue, a deep generative model that employs probabilistic representation of single-nucleus RNA sequencing data to reconstruct the gene expression signals that are lost during nuclei isolation procedures (Figure 1B). The specific method is described in the Methods section. Briefly, the model utilizes a variational autoencoder (VAE) framework to transform the raw, unnormalized count data by modeling gene expression patterns according to a zero-inflated binomial distribution^8,10,12,13^. CytoRescue outputs raw read counts as its prediction target, facilitating seamless integration into existing analytical pipelines. To leverage available paired single-nucleus RNA-seq and single-cell RNA-seq datasets, we implemented CytoRescue as a guided VAE architecture that enables the model to learn from established distributions (Figure 1B)^14^. We curated matched snRNA-seq and scRNA-seq datasets from published literature to construct our training, testing, and validation datasets (Table S1)^2–4^. RM101 was selected as the validation dataset due to the absence of any other data from the same biological sample in the training set (Table S1). Additionally, 8-CST-0892 was randomly selected as validation dataset as well^3,4^.

To train CytoRescue, we first aligned snRNA-seq data with paired scRNA-seq data from identical samples using scmap for cell mapping (Figure S1)^15^. The model employs raw read counts from scRNA-seq as prediction targets to guide the learning of gene expression patterns in snRNA-seq data and recover signals from cytoplasmic content lost during nuclei isolation. Previous studies have demonstrated that microglia activation signals are predominantly lost in snRNA-seq data, as genes associated with microglia activation are enriched in the cytoplasm^4^. We compared pseudobulk gene expression profiles between snRNA-seq and paired scRNA-seq data in microglia from sample RM77 (training dataset) and RM101 (validation dataset) (Figure 1C, 1D). Overall, snRNA-seq data exhibited strong correlation with scRNA-seq data, with Pearson’s correlation coefficients (PCC) being approximately 0.85 (Figure 1C-E). However, disease-associated microglia (DAM) genes were significantly enriched in those genes having higher expression in scRNA-seq than those in snRNA-seq (Figure 1C, 1D). Notably, the majority of DAM genes exhibited greater than 2-fold expression differences, supporting their higher expression in scRNA-seq (sample RM77: 36 vs. 0; sample RM101: 41 vs. 0). This observation is consistent with a previous finding^4^. This trend of a disproportionately high number of genes with elevated expression in scRNA-seq, but not in snRNA-seq, was also observed in lung samples (Figure 1E). We subsequently applied CytoRescue to snRNA-seq data and compared the results with paired scRNA-seq data across all training and validation samples (Figure 1F-H, Figure S3, S4). The rescued gene expression profiles demonstrated significantly improved alignment with scRNA-seq data, with PCC values being increased from 0.894 to 0.971 in sample RM77_2, and from 0.882 to 0.955 in RM101_2, and from 0.825 to 0.915 in 8-CST-0892 (Figure 1F-H). Consistent improvements were observed across all other samples. The uniformly enhanced PCC values in both training and testing datasets indicate that CytoRescue can effectively recover the lost counts in snRNA-seq experiments (Figure 1P).

Given the availability of numerous imputation and denoising tools for single-cell/single-nucleus sequencing data, we investigated whether the lost signals in snRNA-seq could be recovered through conventional imputation or denoising approaches. We selected two widely used tools for comparison: scVI^16^ (based on VAE) and ALRA^17^ (based on low-rank approximation) (Figure 1L-P). As demonstrated, these imputation and denoising tools were not designed to restore discarded cytoplasmic mRNAs from snRNA-seq data and failed to effectively recover the lost signals (Figure 1L-P). Furthermore, we evaluated the consistency of cell type annotations between raw snRNA-seq counts and CytoRescue-predicted counts using adjusted Rand index (ARI), normalized mutual information (NMI), and proportion of consistency metrics^18,19^ (see Methods). Our results demonstrate that while effectively recovering cytoplasmic dynamics, CytoRescue-predicted counts preserve cell type signatures well (Figure 1P, Table S2).

We further investigated the extent to which biological structures were captured by the model and preserved in the output. A standard approach for this assessment involves the use of Uniform Manifold Approximation and Projection (UMAP) for dimensionality reduction and cell clustering, combined with cell subpopulation annotation based on prior knowledge^20–22^. When compared with cell clustering using original read counts (Figure 2A-C), UMAP visualization of predicted read counts effectively preserved cell cluster structure (Figure 2D-F), particularly when it was stratified by sample type (Figure 2A, 2D). When clustered by original read count, microglia cells from brain tissue were intermixed with other immune cells from normal lung and tumor samples (Figure 2B). Following CytoRescue prediction, microglia cells were appropriately separated from other immune cell types as expected (Figure 2E). Additionally, CytoRescue was designed in a raw-in-raw-out manner, (Figure 2C, 2F), which facilitates seamless integration into existing analytical pipelines. We extended our analysis to examine the latent space representation to further verify whether CytoRescue can capture the underlying biological features in snRNA-seq data, but not just memorizing the training data^7^. As demonstrated in Figure 2G-I, the cells were well-clustered and appropriately separated regardless sample types, cell types, and patients we examined, with the most distinct separation being observed when stratified by sample type (Figure 2G). In summary, our evaluation demonstrated that CytoRescue could effectively capture biological structures in snRNA-seq data and recover the signals from discarded cytoplasmic content while preserving biological signatures.

**Figure 2.**
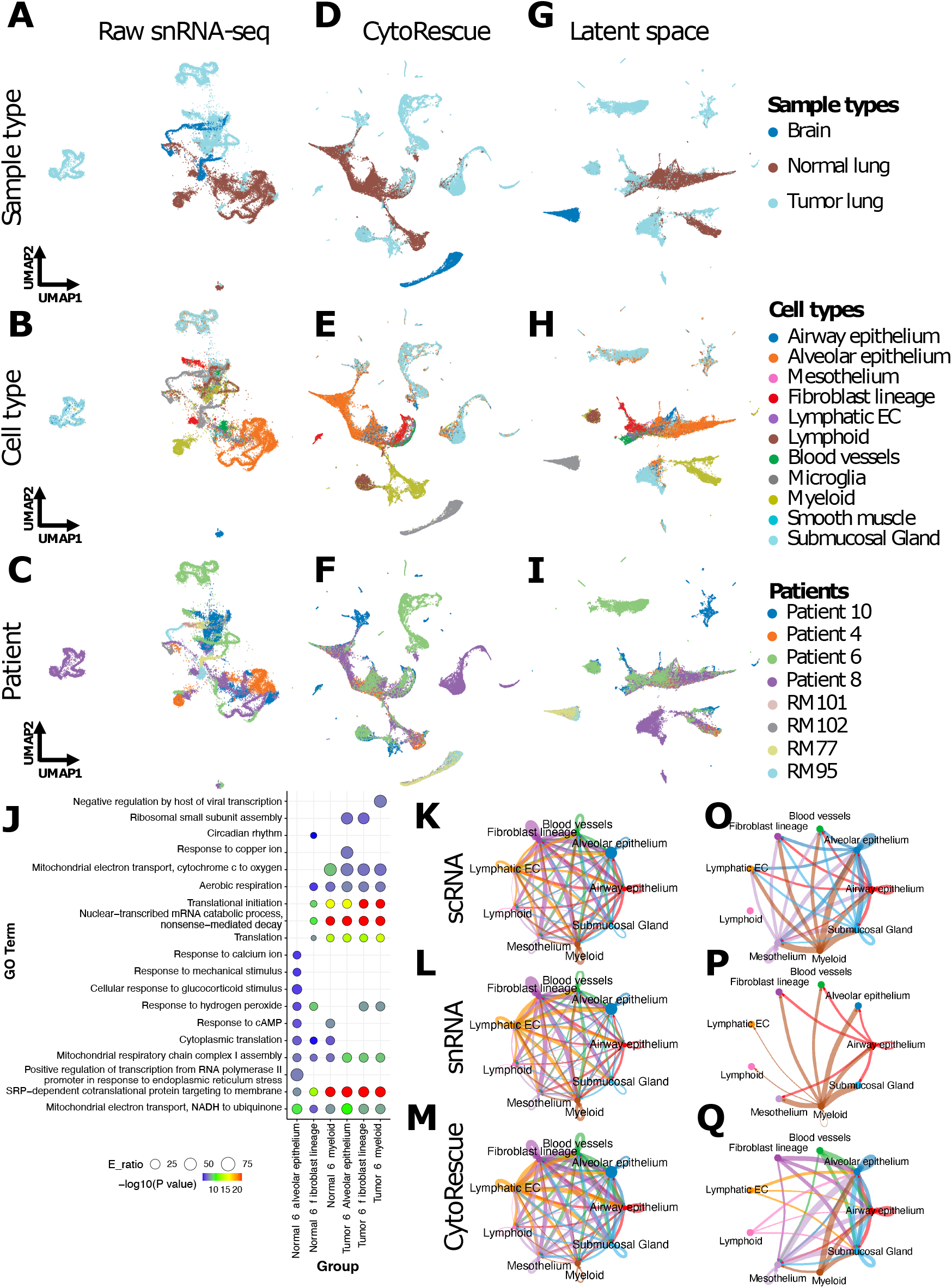
CytoRescue preserves biological features in original data and recovers cytoplasmic dynamics. Uniform Manifold Approximation and Projection (UMAP) of all cells in this study using raw snRNA-seq counts (A-C), CytoRescue-predicted counts (D-F), and latent representations from CytoRescue (G-I). Cells are colored by sample type (A, D, G), cell type (B, E, H), and patient ID (C, F, I) to examine different aspects of biological structure. (J) Gene Ontology Biological Process terms enriched in the genes with greater than 2-fold expression change in CytoRescue-predicted profiles compared to raw snRNA-seq counts. Dot size represents enrichment ratio; color represents significance (−log_□ □_(P value)) as determined by Fisher’s exact test. (K-M) Cell-cell communication networks identified from validation sample 8-CST-0892. Interaction weights across all signaling pathways are illustrated for scRNA-seq counts (K), snRNA-seq counts (L), and CytoRescue-predicted counts (M). (O-Q) Cell-cell communication interactions through the EGF signaling pathway identified from scRNA-seq counts (O), snRNA-seq counts (P), and CytoRescue-predicted counts (Q).

We compared the expression profiles before and after the CytoRescue prediction. This comparison identified the genes exhibiting greater than 2-fold increased expression in tumor and normal lung samples across three cell types with more than 100 cells present in all analyzed samples (Figure S5). For these differentially expressed genes, we performed Gene Ontology (GO) based gene set enrichment analysis to identify overrepresented biological processes^23^. As anticipated, the most significantly enriched pathways across samples were associated with translation, mitochondrial energy production, and response to external stimuli (Figure 2J). Notably, the primary differences in enriched pathways among distinct cell types and samples were related to external stimulation responses, which is consistent with the diverse biological contexts of different cell types and sample conditions (Figure 2J, Figure S5). Previous studies have demonstrated that cell-cell communications (CCC), particularly those mediated by ligand-receptor (L-R) interactions, are essential for coordinating multiple cell populations in lung tissue to regulate responses to stimuli^24,25^. To evaluate whether CytoRescue can restore CCC signals, we analyzed the validation lung sample 8-CST-0892, which was withheld from training. Following the standard CellChat pipeline, we inferred CCC between major lung cell types using raw scRNA-seq counts, snRNA-seq counts, and CytoRescue-predicted counts (Figure 2K-Q)^26^. Compared to CCC inferred from snRNA-seq counts, scRNA-seq data yielded significantly higher numbers of interactions (Figure 2K-M), consistent with previous findings^3^. Notably, previous studies have shown that epithelial cells dominate in the EGF-based L-R interaction networks^27^. However, interactions between alveolar epithelium and other cell types were predominantly lost in snRNA-seq data (Figure 2P), whereas these signals were successfully identified in both scRNA-seq data and CytoRescue-predicted counts (Figure 2O, 2Q). Collectively, CytoRescue successfully recovered essential biological signals in a dataset never encountered during training.

## Conclusions

We present CytoRescue, a generative AI model designed to recover lost signals in single-nucleus RNA-sequencing experiments. We demonstrate that CytoRescue effectively restores lost cytoplasmic dynamics during nuclei isolation while preserving biological structures in both latent space and output representations. Additionally, CytoRescue employs a raw-in-raw-out architecture that enables seamless integration into existing single-cell RNA-seq analytical pipelines. When applied to the samples excluded from our training, CytoRescue could successfully restore cell-cell communication signals between alveolar epithelium and other cell types. While more evaluation work based on paired scRNA-seq and snRNA-seq datasets is needed, CytoRescue is demonstrate useful for detecting and evaluation of cytoplasmic gene expression signals from snRNA-seq data.

## Methods

### Paired scRNA-seq and snRNA-seq data sets

Paired scRNA-seq and snRNA-seq datasets from brain and lung tissues were utilized in this study^3,4^. For brain samples, raw count data from single microglia cells were obtained from Gene Expression Omnibus (GEO) datatset GSE137444 (scRNA-seq) and GSE153807 (snRNA-seq)^4^. For lung samples, paired scRNA-seq and snRNA-seq count data from normal and tumor lung tissues were obtained from Zenodo (accession: 10.5281/zenodo.1120562)^3^.

### Cell type annotation and mapping

Cell type annotation for the collected datasets was performed following a standard reference-based pipeline implemented in Seurat (v5.0.0)^28^. For brain snRNA-seq data, cell types were annotated by anchoring to a reference subsampled to a maximum of 2,000 cells per cell type from the SEA-AD reference^29^ For lung samples, cell types were annotated using a reference subsampled to a maximum of 2,000 cells per cell type from HLCA (v1.0)^30^. Following annotation, cells in snRNA-seq data were mapped to corresponding cells in paired scRNA-seq data using scmap^15^.

### Model structure and training

In VAE architectures, Gaussian distributions are commonly employed due to their differentiability during reparameterization^22^. Here, we designed the latent representation *z* to be Gaussian distribution, and the zero-inflated binomial parameters were inferred from input and output layers, effectively implemented ZINB modeling of single cell/nuclei data^6,22^.

We first define negative binomial probability mass function as:

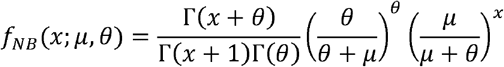

To guide the model to learn from matched snRNA-seq and scRNA-seq data, ZINB distribution is defined: *x* denotes expression data, *μ* as mean, *θ* as the dispersion parameter and *π* as zero-inflation probability. We designed the loss function as:

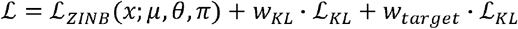

Where ℒ_ziNB_ denotes the ZINB loss of observed count x, which is defined by:

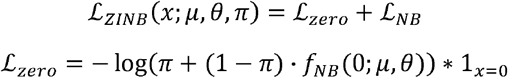

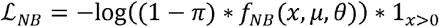

where 1<_*x*-0_ is the indicator function equals to 1 when x=0, otherwise 0. _1x>0_ is the indicator function equals to 1 when x>0, otherwise 0. *W*_*KL*_is the loss weight parameter, while *ℒ*_*KL*_ denotes the Kullback-Leibler (KL) divergence loss measured between approximate posterior (defined by *μ* as mean, and *σ* as variance) and standard Gaussian distribution in latent space by:

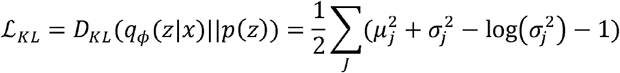

Meanwhile W_target_ is the loss weight parameter of target loss ℒ_KL_, which is difined by:

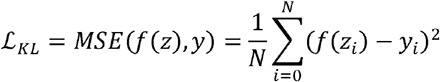

For model training, we held out 2 samples from training as validation dataset for independent test. Further, the training datasets were shuffled and randomly selected 80% as training data while 20% as testing data. The model is trained on Ubuntu server equipped with 1 AMD EPYC 7742 64-Core Processor and 8 NVIDIA A100-SXM4-40GB GPUs, only 1 GPU was used during training and prediction.

### Evaluation

We evaluated how CytoRescue recovers cytoplasmic gene expression dynamics while preserving biological structures in training, testing, and validation datasets through: 1) assessing concordance between predicted counts and paired scRNA-seq data, 2) comparing concordance between counts processed by denoising/imputation tools and paired scRNA-seq data, 3) evaluating clustering consistency between snRNA-seq counts and CytoRescue-predicted counts, and 4) comparing dimensionality reduction maps of all cells across raw snRNA-seq, scRNA-seq, and predicted count data.

The consistency between different processing procedures is measured by Pearson’s correlation of the mean expression for each gene as:

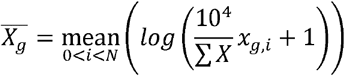

where *X* denotes the gene expression matrix in each sample.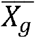 denotes mean expression of a specific gene *g,N* is the total number of cells.

The clustering consistence is measured by adjusted Rand index (ARI)^19^. ARI, which is normalized mutual information (NMI)^18^ and proportion of consistent (Consistency) label between unprocessed and predicted profiles, is calculated as follows:

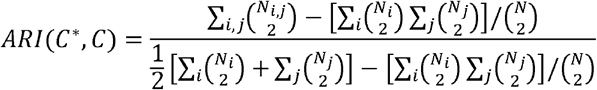

Where *C*^*^denotes the cell type annotation from CytoRescue’s predicted counts and *C* denotes the cell type annotation from raw snRNA-seq counts. N denotes total number of data points. *N*_i,j_denotes number of data points labeled as both cell types *i* in *C*^*^ and *j* in *C. N*_*i*_ denotes number of data points labeled as cell type *j* in .*C*

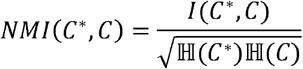

where

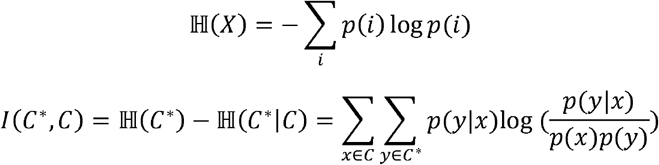

Then, we calculate

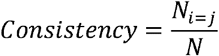

where *N*_*i=j*_ denotes number of cells with the same cell type in *C*^*\**^ and *C*. And N denotes total number of data points in *C*^*\**^.

The dimension reduction of cells was done by umap-learn package (v0.5.7) on log normalized read counts for each group.

### Gene set enrichment analysis

The most recent Gene Ontology annotation and mapping files were downloaded from the Gene Ontology Consortium (GOC)^23^. Biological Process (BP) terms were used for pathway enrichment analysis of differentially expressed genes. Fisher’s exact test was employed to identify significantly overrepresented GO terms.

### Cell-cell communication analysis

Raw read counts from scRNA-seq, snRNA-seq, and CytoRescue-processed data were normalized to a total count of 10,000 per cell and subsequently log-transformed. CellChat (v2) was then applied to the normalized read counts following the standard pipeline with all signaling pathways^26^. The number of connections and contribution of ligand-receptor (L-R) pairs in each circle plot were also inferred.

## Code availability

The source code of CytoRescue and analysis code involved in this paper is deposited at GitHub and can be accessed.

## Data availability

All the data are publicly available as we described in the Methods section.

## Declarations

Competing interests Non declared.

## Acknowledgement

We thank Dr. Xiaoqian Jiang for allowing us to access to GPU resources.

## Supplementary Figures and Tables

**Figure S1**. Data processing workflow in this study. The snRNA-seq data retrieved from the literature were processed using standard filtering and cell type annotation procedures. All cell types from lung samples and microglia cells from brain samples were retained for scMAP mapping to paired scRNA-seq data. A snRNA-to-scRNA paired dataset was ultimately prepared for training, testing, and analysis.

**Figure S2**. Training process of CytoRescue. Training was terminated at 30 epochs when both training and validation losses converged after 25 epochs.

**Figure S3**. Log-normalized mean gene expression comparison between scRNA-seq and snRNA-seq, CytoRescue-predicted counts, scVI-imputed counts, and ALRA-imputed counts in lung samples.

**Figure S4**. Log-normalized mean gene expression comparison between scRNA-seq and snRNA-seq, CytoRescue-predicted counts, scVI-imputed counts, and ALRA-imputed counts in brain samples.

**Figure S5**. Gene Ontology Biological Process terms enriched in the genes with greater than 2-fold change in mean expression in CytoRescue-predicted counts compared to snRNA-seq counts in samples from (A) patient #8, and (B) patient #10.

**Figure S6**. *ERBB2* expression in (A) scRNA-seq, (B) snRNA-seq, and (C) CytoRescue-predicted counts. Expression values are log_10_-normalized.

**Table S1**. Statistics and summary of training and validation datasets in this study.

**Table S2**. Pearson’s correlation coefficients of mean gene expression between scRNA-seq and snRNA-seq by three methods: CytoRescue-predicted, scVI-imputed, and ALRA-imputed counts.

