## Supplementary figures and images for "Cytoplasmic dynamics are overlooked in single nuclei RNA-seq but can be rescued by CytoRescue, a generative AI model to recover cytoplasm enriched gene"

### Figure S1

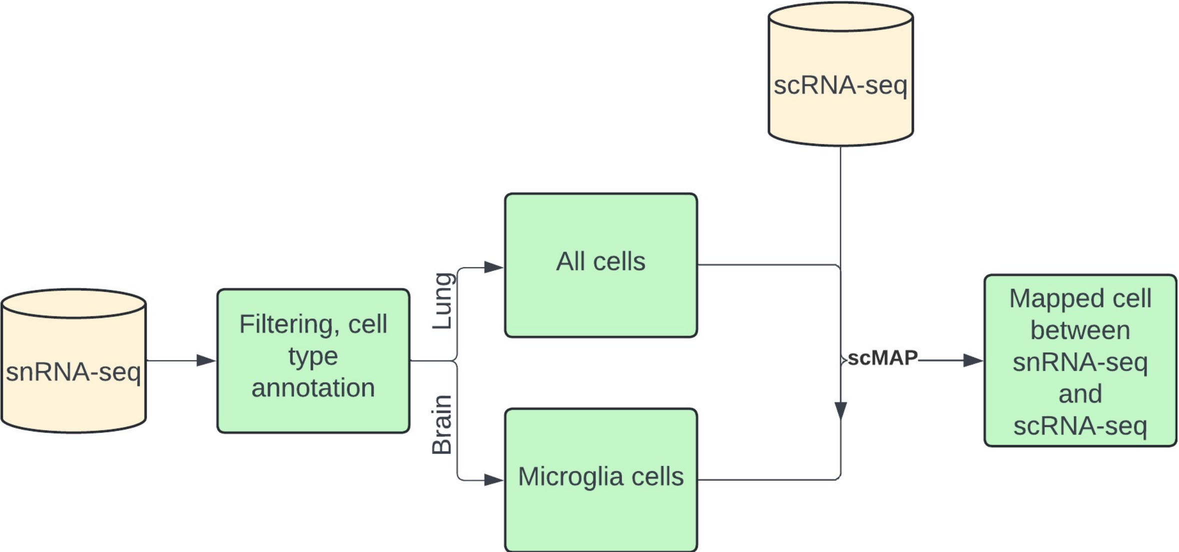

### Figure S2

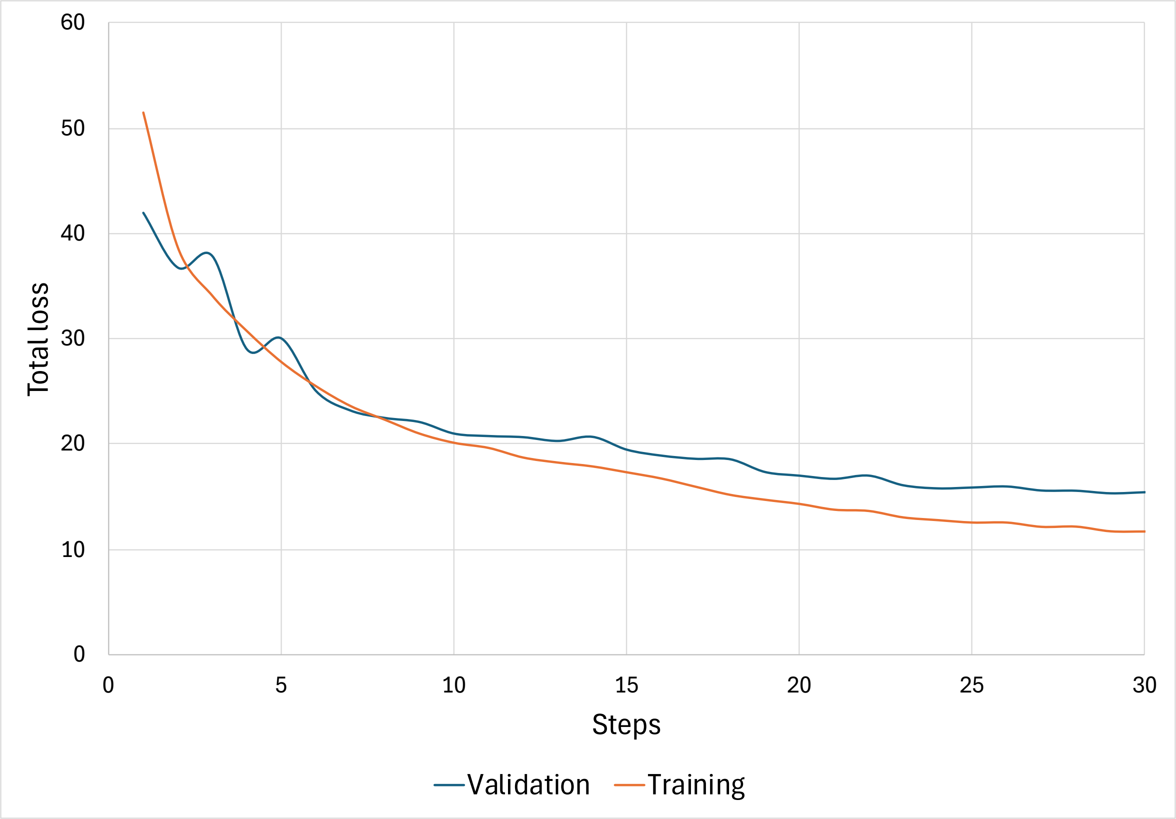

### Figure S3

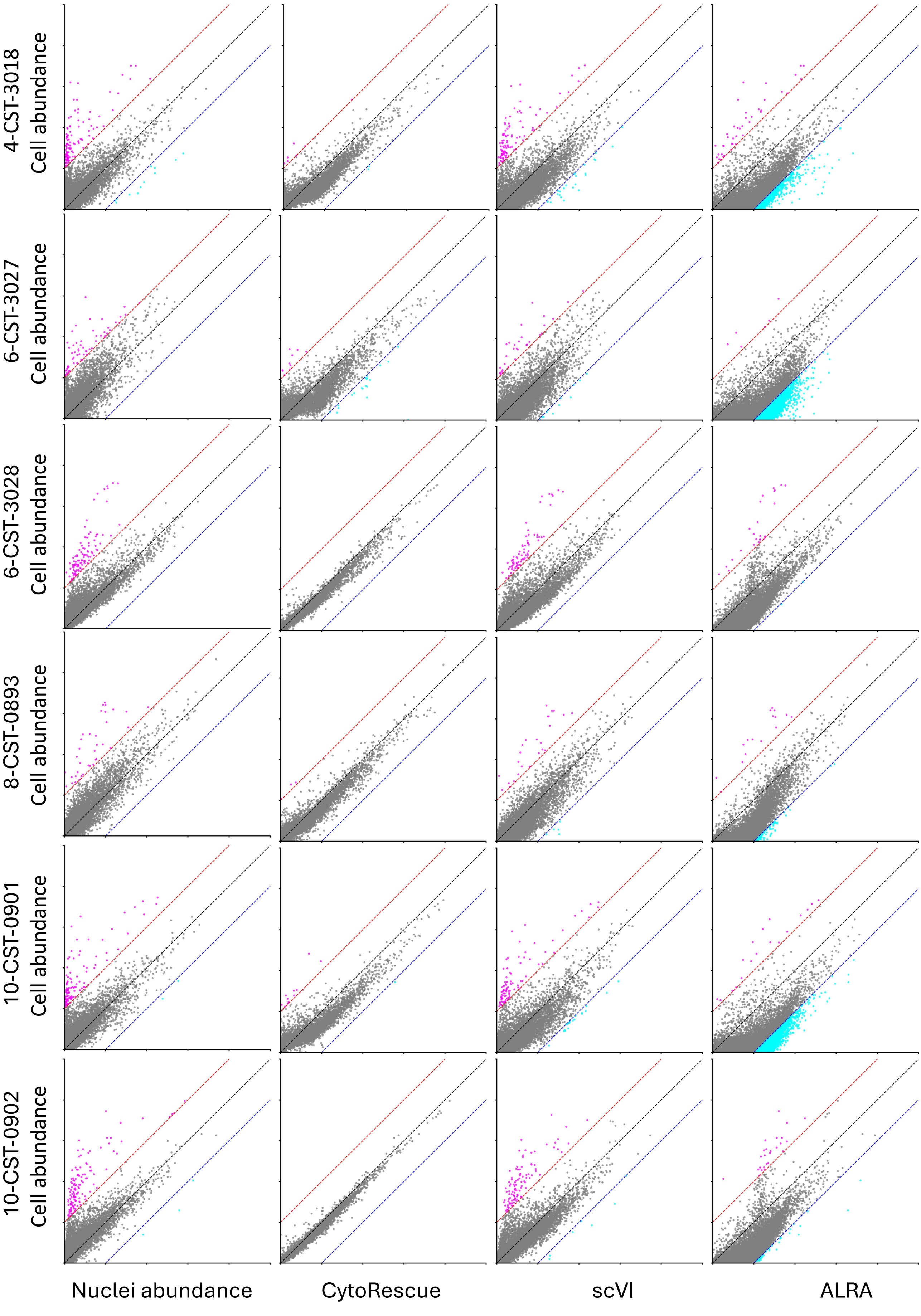

### Figure S4

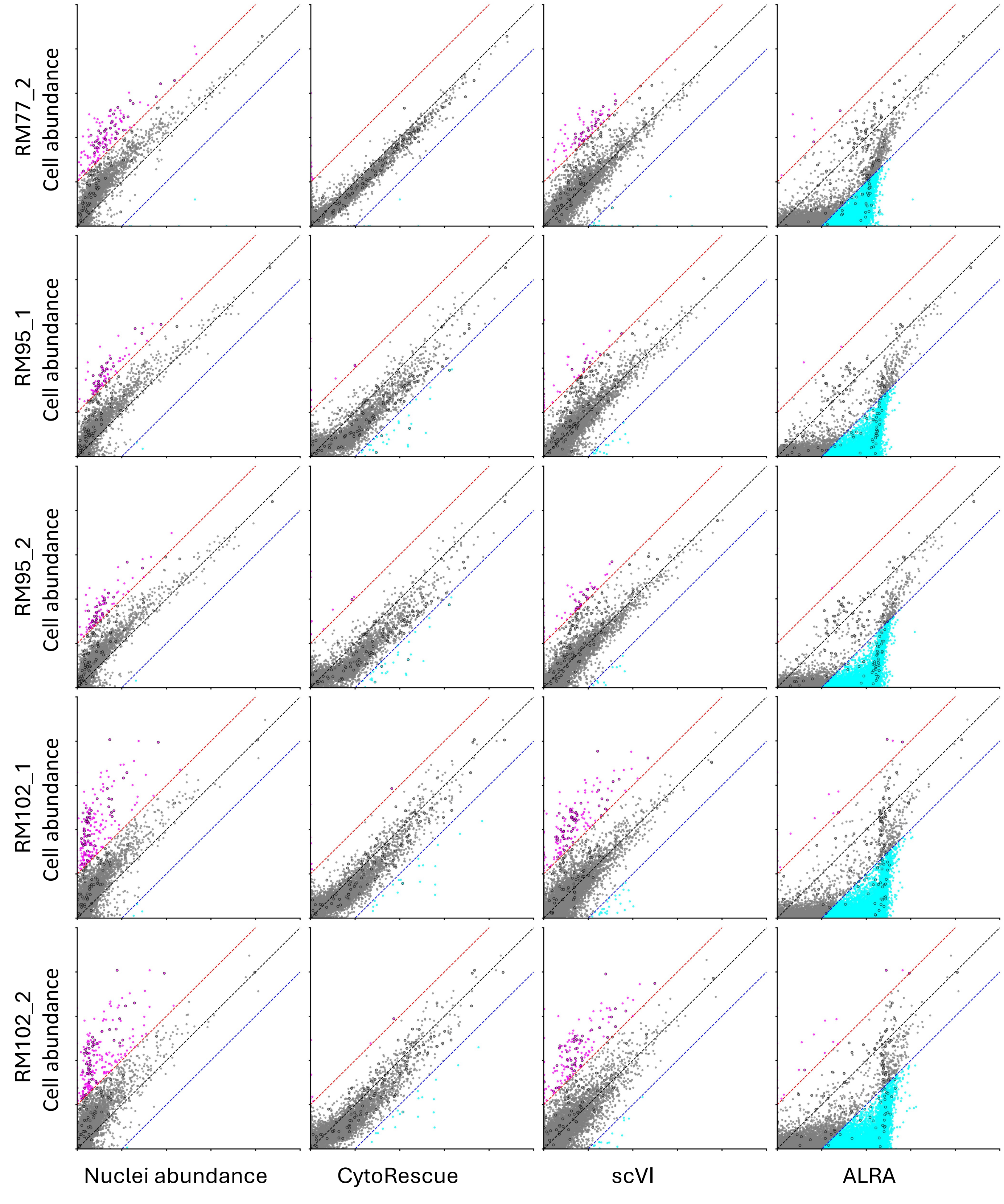

### Figure S5

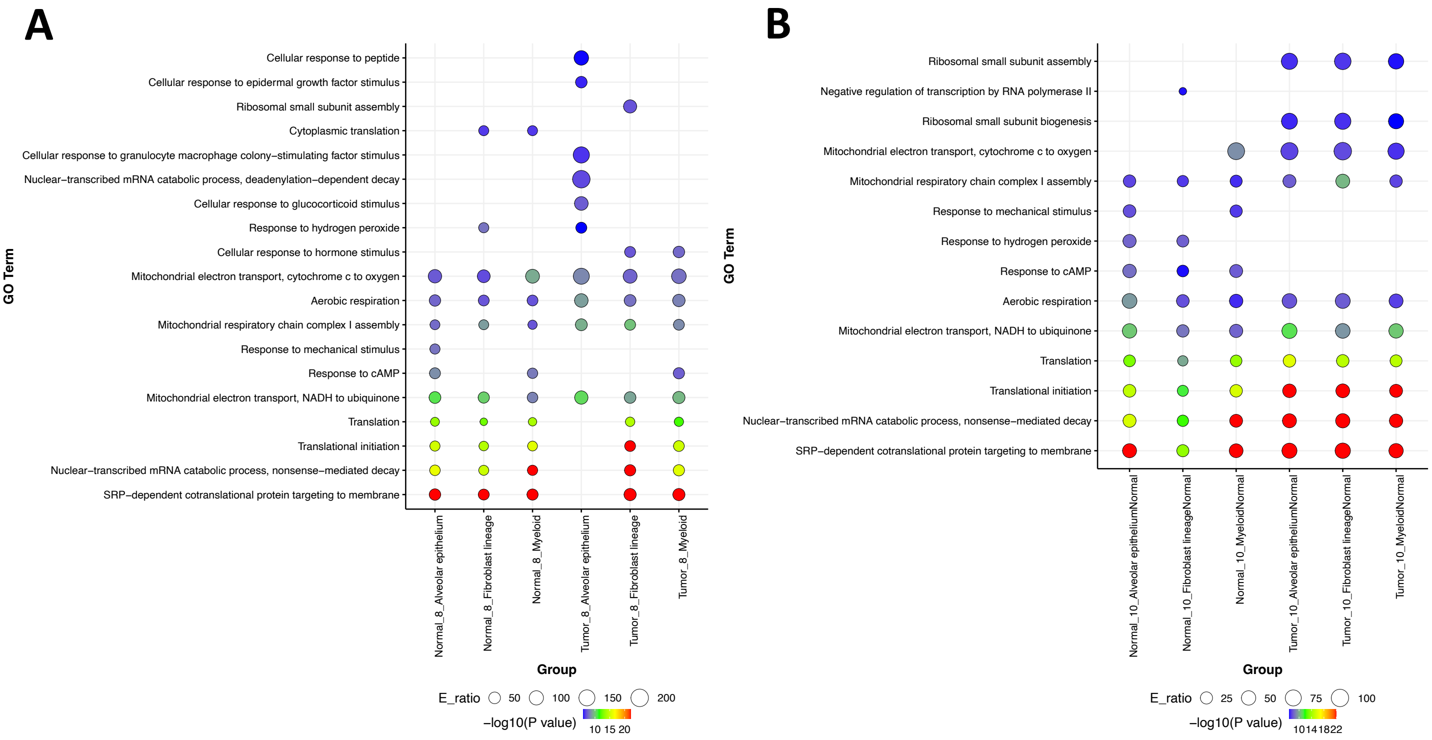

### Figure S6

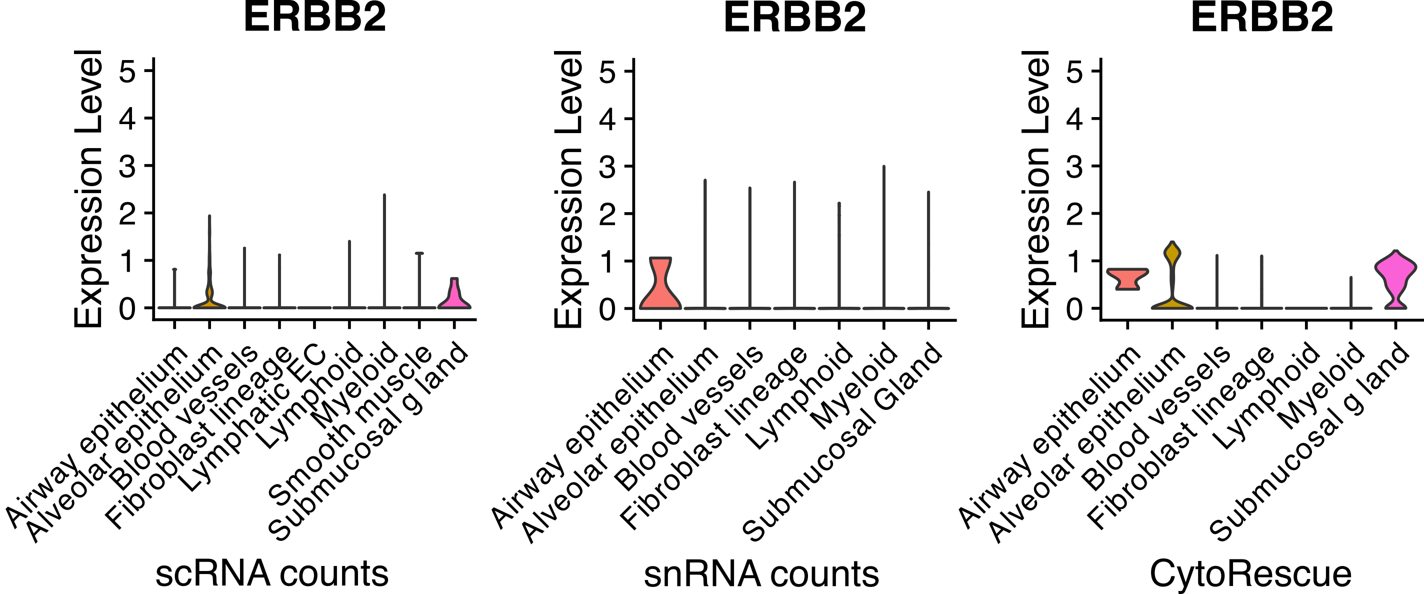
